# SIRPα sequesters SHP-2 to promote IL-4/13 signaling and alternative activation of macrophages

**DOI:** 10.1101/2021.08.06.455421

**Authors:** Lei Shi, Koby Kidder, Zhen Bian, Samantha Kuon Ting Chiang, Corbett Ouellette, Yuan Liu

**Affiliations:** Program of Immunology & Molecular Cellular Biology, Department of Biology, Georgia State University, Atlanta, GA 30302; Center of Diagnostics and Therapeutics, Georgia State University, Atlanta, GA 30302

## Abstract

The Th2 cytokines IL-4 and IL-13 through activation of their shared receptor IL-4Rα direct macrophage alternative activation to promote immunosuppression and wound healing. However, the mechanisms that control macrophage responses to IL-4/13 are not fully understood. Apart from driving JAK-STAT and PI3K-Akt pathways to polarize macrophages toward the alternative phenotype, the activated IL-4/13 receptors recruit negative regulators SHP-1 and SHP-2, which dephosphorylate IL-4Rα and decrease its signaling. Here we report that SIRPα spatially restricts SHP-2 and, by such, promotes IL-4/13 signaling and macrophage alternative activation. SIRPα executes this regulation via its cytoplasmic ITIMs/ITSMs that undergo phosphorylation by IL-4/13-induced, Src kinase-activated Bruton’s tyrosine kinase (Btk), resulting in recruitment of SHP-2 and preclusion of SHP-2 from binding to and inhibiting IL-4/13 receptors. Despite that this regulation occurs independent of CD47, extracellular CD47 ligation of SIRPα facilitates its cytoplasmic phosphorylation and SHP-2 sequestration, leading to stronger IL-4/13 signaling and enhanced macrophage expression of IL-10, TGFβ, CD206, arginase-1, etc. Conversely, deficiency of SIRPα allows SHP-2 to freely bind to γC or IL-13Rα1 and through which dephosphorylate IL-4Rα, dampening its signaling. Consistent with these findings, impaired wound healing in Sirpα^−/−^ mice under experimental colitis correlated with a deficit of immunosuppressive macrophages in the colon, a condition that was corrected by transfusion of ex vivo-produced SIRPα^high^ alternatively activated macrophages.

## Introduction

Alternative activation (M2/M2a) of macrophages by the Th2 cytokines IL-4 and/or IL-13 plays an important role in macrophage-driven clearance of parasitic infection, resolution of inflammation and wound healing, as well as dysregulated pathological conditions including allergy, excessive fibrosis and immunosuppressive milieus conducive to tumor growth and metastasis (*1, 2*). To drive the typical M2a phenotype, IL-4 and IL-13 bind to their cognate receptor chain, IL-4Rα and IL-13Rα1, respectively, and induce receptor and co-receptor dimerization forming IL-4Rα/γC and/or IL-4Rα/IL-13Rα1 heterodimers, which enable activation of their associated Janus kinases (JAK1/3, Tyk2) to phosphorylate the receptor cytoplasmic domains for the docking of downstream signaling molecules including STAT6, Gab2 and insulin receptor substrate (IRS2) (*3–5*). After these signaling molecules dock to cytokine receptors, JAKs/Tyk2 proximally phosphorylate them and lead to their activation to mediate downstream effector functions. For example, activated STAT6 undergoes dimerization and then translocates to the nucleus to induce critical gene transcription, conferring macrophage alternative activation and immunosuppressive functions (*3, 4*). Phosphorylated Gab2 and IRS2 initiate scaffolding of downstream molecules, leading to PI3K-Akt1/2 and other pathways that also promote immunosuppressive macrophage polarization (*5, 6*). However, the phosphorylated IL-4/13 receptors recruit not only signaling molecules potentiating an alternative phenotype, but also SH2 domain-containing tyrosine phosphatases SHP-1 and SHP-2, which dephosphorylate the receptors and temper IL-4/13 signaling. Indeed, Tachdjian et al. have shown that preventing SHP-1 from docking to IL-4Rα (Y709) enhances IL-4/13 signaling and exacerbates allergy-associated pulmonary inflammation (*7*). Other studies have also shown that SHP-2 association with the IL-4 receptor suppresses macrophage alternative polarization and that inhibition of SHP-2 by a specific inhibitor PHPS1 or SHP-2 deficiency predisposes macrophages toward an anti-inflammatory phenotype (*8–10*).

SIRPα is a myeloid leukocyte-expressed immunoreceptor tyrosine-based inhibitory/switch motifs (ITIMs/ITSMs)-containing signaling receptor whose canonical function, via interacting with the self-marker CD47, is to inhibit professional phagocytes, e.g. macrophages and dendritic cells, from phagocytosing self-cells. Besides phagocytosis, SIRPα also inhibits macrophage and granulocyte inflammatory responses and dampens their proinflammatory cytokine and ROS production (*11–15*). Under immunosuppressive conditions, our previous study found that SIRPα mediates different regulation, which instead of executing inhibition, promotes macrophage alternative activation to display an anti-inflammatory phenotype (*16, 17*). The detailed mechanisms by which SIRPα mediates distinctive regulations under varied stimulations remain unresolved, except that the phosphorylation of SIRPα cytoplasmic ITIMs/ITSMs and their recruitment of SHP-1 and/or SHP-2 are likely involved (*14, 16, 18, 19*). However, neither the stimulatory state under which SIRPα recruits either SHP-1 or SHP-2 nor the regulatory ramifications arising from differential SHP-1 or SHP-2 binding has been clearly demonstrated.

In the present study, we report that upon IL-4 and IL-13 stimulation, SIRPα mediates a ‘dis-inhibition’ mechanism that, through sequestering SHP-2, promotes macrophage polarization toward an immunosuppressive, alternative activation phenotype. This regulation is initiated by an IL-4/13-induced and Src kinase-activated Bruton’s tyrosine kinase (Btk), which phosphorylates the cytoplasmic tyrosines within SIRPα ITIMs/ITSMs in a manner that leads to exclusive docking of SHP-2, but not SHP-1. By recruiting SHP-2, SIRPα depletes SHP-2 from accessing IL-4/13 receptors (γC and IL-13Rα1 specifically), thereby enhancing IL-4Rα phosphorylation and downstream activation of STAT6, Gab2 and PI3K-Akt. Consistent with these findings, deficiency of SIRPα allows SHP-2 to freely inhibit IL-4/13 receptors and subsequently dampen macrophage alternative activation, a condition associated with hindered resolution of inflammation in a murine colitis-recovery model. Converesly, transfusion of alternatively activated macrophages expressing increased SIRPα restored the anti-inflammatory response, promoting post-colitis wound healing and recovery.

## Results

### SIRPα promotes IL-4- and IL-13-induced macrophage alternative activation

To test the effect of SIRPα on IL-4-induced alternative activation, murine macrophages freshly isolated from the peritoneum (PEM) and derived from the bone marrow (BMDM) with and without SIRPα expression (WT and Sirpα^−/−^, respectively) were treated with IL-4 in the presence or absence of mCD47.ex, a soluble murine CD47 extracellular domain that ligates SIRPα on macrophages (*11, 20*). The same experiments were also done with human macrophages (HMM) derived from peripheral blood monocytes in the presence or absence of a human CD47 extracellular domain (hCD47.ex) (*19, 21*). Both murine and human macrophages treated with IL-4 produced significantly higher IL-10 and TGFβ when their SIRPα was ligated by CD47 (mCD47.ex or hCD47.ex) (Fig. 1, A-C). In contrast, SIRPα deficiency (Sirpα^−/−^) blunted IL-4 responsiveness and markedly reduced IL-10 and TGFβ production. As expected, Sirpα^−/−^ macrophages were not affected by the presence of CD47 ligation. Aside from impacting cytokine production, CD47 ligation of SIRPα also enhanced the expression of CD206, arginase-1 (Arg-1) and the transcription of Arg1, Ym1, Msr2 and Fizz1 in IL-4-treated WT murine macrophages, whereas SIRPα deficiency largely reduced these phenotypic markers (Fig. 1, D-F). Investigation of IL-4 downstream signaling confirmed that CD47-SIRPα ligation bolstered the activation of STAT6 and PI3K-Akt pathways, having increased phosphorylation of STAT6 (p-STAT6), Akt1 and Akt2 (p-Akt1 and p-Akt2), whereas SIRPα deficiency dampened IL-4-induced phosphorylation of these molecules (Fig. 1G). Similar SIRPα-mediated regulation of IL-4 signaling was also observed in bone marrow-derived dendritic cells (BMDC) with the presence or absence of SIRPα respectively promoting or diminishing IL-4 signal transduction (Fig. 1H).

**Figure 1.**
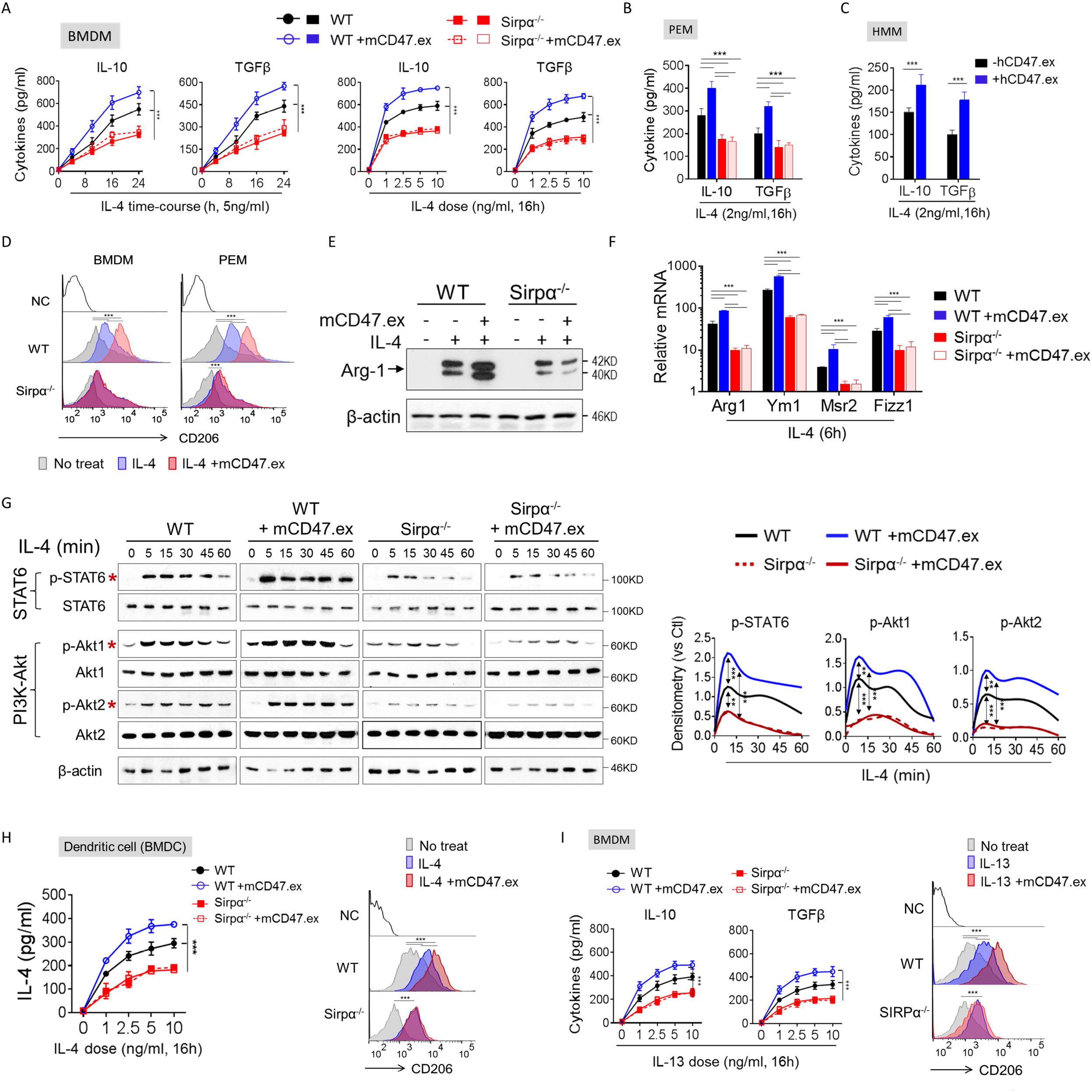
SIRPα promotes IL-4/13-induced macrophage alternative activation. (**A**) WT and Sirpα^−/−^ BMDM were treated with IL-4 for indicated time periods or by various doses of IL-4 in the presence or absence of mCD47.ex, and the levels of IL-10 and TGFβ in the supernatants were tested by ELISA. (**B**) PEM from WT and Sirpα^−/−^ mice were also activated by IL-4 (2ng/ml) in the presence or absence of mCD47.ex, followed by analysis of IL-10 and TGFβ secretion after 16h. (**C**) The production of IL-10 and TGFβ by IL-4-treated human PBMC-derived macrophages (HMM) in the presence or absence of hCD47.ex. (**D**) The expression of CD206 in WT and Sirpα^−/−^ macrophages detected by flow cytometry. (**E**) The level of Arg-1 in WT and Sirpα^−/−^ BMDM determined by WB. (**F**) The transcription of Arg1, Ym1, Msr2 and Fizz1 in WT and Sirpα^−/−^ BMDM examined by RT-PCR. (**G**) The phosphorylation of STAT6, Akt1 and Akt2 in WT and Sirpα^−/−^ BMDM during IL-4-induced activation. The phosphorylation of STAT6, Akt1 and Akt2 was quantified by densitometric analysis and normalized against the corresponding total protein. (**H**) WT and Sirpα^−/−^ BMDC were treated with various doses of IL-4 for 2h in the presence or absence of mCD47.ex. The levels of biologically active (released) IL-4 in fresh complete medium were tested by ELISA and the expression of CD206 in IL-4 (10ng/ml)-treated WT and Sirpα^−/−^ BMDC after 16 h (*49, 53*). (**I**) The production of IL-10 and TGFβ by various doses of IL-13-treated BMDM in the presence or absence of mCD47.ex and the expression of CD206 in IL-13 (10ng/ml)-treated WT and Sirpα^−/−^ macrophages detected by flow cytometry. Data are presented as mean ± SEM of at least three independent experiments (n ≥ 3). Statistical significance was determined by two-way ANOVA with Tukey’s post hoc test (**A**, **G**, **H**, **I**) or Student’s t test (**B**, **C**, **D**, **F**, **H**, **I**), where **p < 0.01 and ***p < 0.001. All blots are representative of at least three independent experiments.

SIRPα similarly enhanced the alternative activation of macrophages stimulated by IL-13, which shares the IL-4 signaling cascade via the IL-4Rα/IL-13Rα1 complex (*22*). As with IL-4 stimulation, SIRPα ligation by CD47 increased the secretion of IL-10 and TGFβ and the expression of CD206 in IL-13-treated macrophages, whereas SIRPα deficiency markedly diminished these IL-13-driven effects (Fig. 1I).

### IL-4/13 induce phosphorylation of SIRPα cytoplasmic domain and recruitment of SHP-2

To determine the mechanism by which SIRPα promotes IL-4/13-induced macrophage alternative activation, we assessed SIRPα cytoplasmic ITIMs/ITSMs phosphorylation and their association with SHP-1 and/or SHP-2. As shown (Fig. 2, A-B), IL-4 stimulation dose-dependently and rapidly (< 5min) induced robust tyrosine phosphorylation in SIRPα (SIRPα^PY^). Moreover, IL-4-driven phosphorylation of SIRPα occurred independent of extracellular ligation by CD47, albeit the latter enhanced the phosphorylation intensity. Without IL-4 stimulation, mere CD47 ligation was inefficient, inducing only weak SIRPα phosphorylation. These results suggest that IL-4 activates a special tyrosine kinase(s), which phosphorylates SIRPα ITIMs/ITSMs, and that CD47 ligation triggers a structural change in the SIRPα cytoplasmic domain that facilitates the tyrosine phosphorylation to occur. Detecting SHP-1 and SHP-2 association with SIRPα by co-immunoprecipitation (IP) found that phosphorylated SIRPα nearly exclusively recruited SHP-2, not SHP-1, which remained in the SIRPα-excluded post-bound fraction (Fig. 2, A-B). Two forms of SHP-2 were detected: one having a higher molecular weight (MW; 75kDa) that was phosphorylated (p-SHP-2 or SHP-2^Tyr542^) and another being non-phosphorylated with lower MW (72kDa, SHP-2). Only phosphorylated SHP-2 bound to SIRPα^PY^ upon IL-4 stimulation (Fig. 2, A-B). SHP-2 phosphorylation was dose-dependently correlated with IL-4 stimulation, independent of SIRPα expression or CD47 ligation (Fig. 2C).

**Figure 2.**
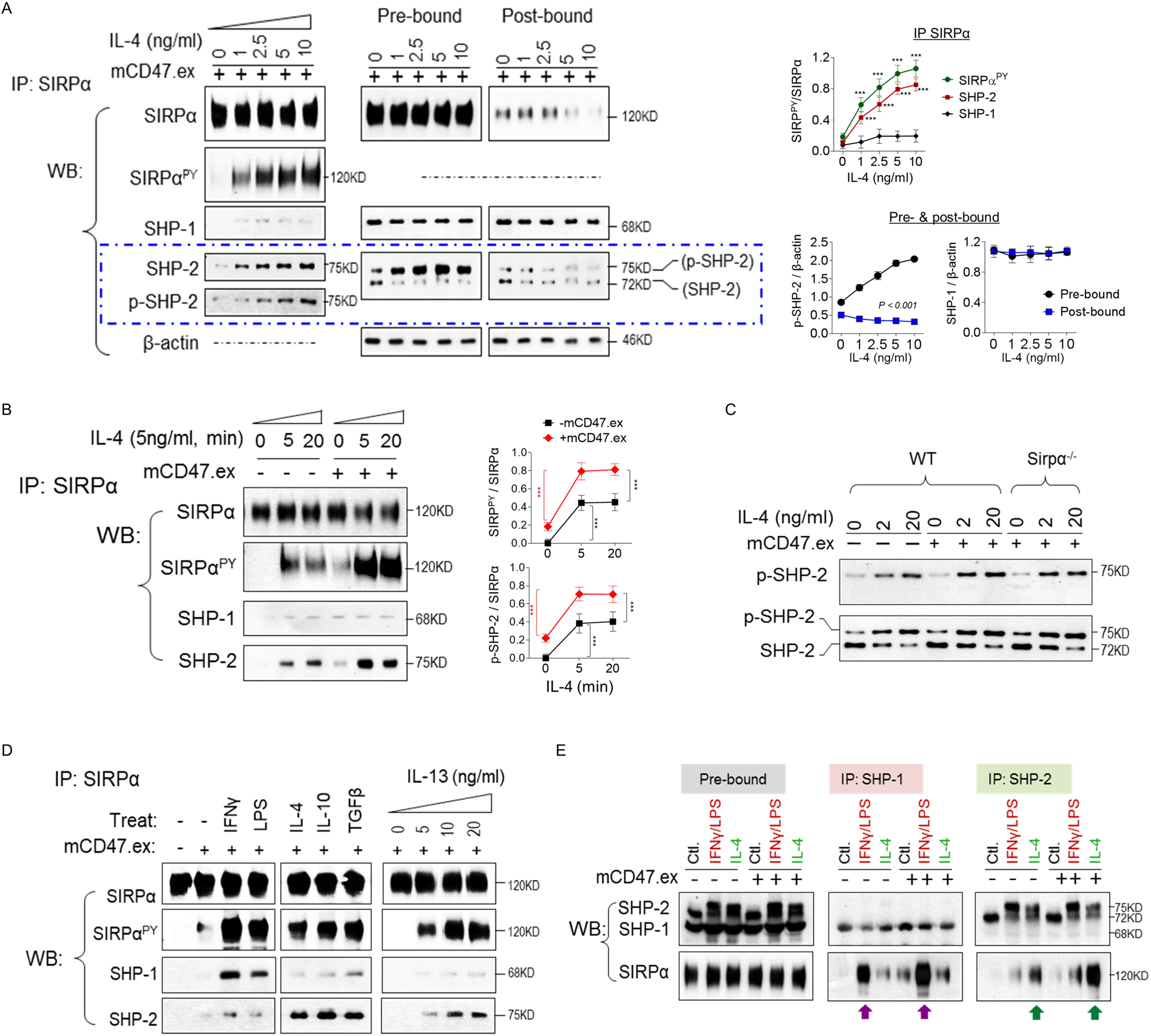
IL-4/13 induces phosphorylation of SIRPα cytoplasmic domain and the recruitment of SHP-2. (**A**) WT BMDM were treated with various doses of IL-4 for 30 min in the presence of mCD47.ex. SIRPα phosphorylation and SHP-1/2 association were assessed following IP of SIRPα. (**B**) WT BMDM were treated with IL-4 (5ng/ml) for 5 or 20min in the presence or absence of mCD47.ex. SIRPα phosphorylation and its binding to SHP-1/2 were assessed following IP of SIRPα. (**C**) Dose-dependent effect of IL-4 on phosphorylation of SHP-2. Note: IL-4 dose-dependently enhances the level of phosphorylation of SHP-2 on Tyr542 and SIRPα signaling has no effect on SHP-2 phosphorylation. (**D**) WT BMDM were treated with IFNγ (20ng/ml), LPS (100ng/ml), IL-4 (5ng/ml), IL-10 (20ng/ml), TGFβ (20ng/ml) or various doses of IL-13 in the presence of mCD47.ex for 20min and the association of SHP-1 and SHP-2 with SIRPα was detected by IP of SIRPα. (**E**) WT BMDM were treated with IFNγ (20ng/ml)/LPS (100ng/ml) or IL-4 (5ng/ml) for 20min and the selective association of SIRPα with SHP-1 or SHP-2 was also detected and quantified by IP of SHP-1 or SHP-2. Meanwhile, macrophages were treated with mCD47.ex to ligate SIRPα during both activations. Data are presented as mean ± SEM of at least three independent experiments (n ≥ 3). Statistical significance was determined by Student’s t test (**A**) or two-way ANOVA with Tukey’s post hoc test (**A**, **B**) where ***p < 0.001. All blots are representative of at least three independent experiments.

Similar to IL-4, IL-13 stimulation induced SIRPα phosphorylation and association of phosphorylated SHP-2, but not SHP-1 (Fig. 2D). SIRPα^PY^ selectively recruiting SHP-2 also occured in macrophages stimulated with IL-10 and TGFβ, whereas macrophages stimulated with proinflammatory (M1) factors IFNγ and LPS comprised phosphorylated SIRPα that instead associated with SHP-1 (Fig. 2D). Immunoprecipitating SHP-1 or SHP-2 confirmed that SHP-1 co-precipitated with SIRPα in IFNγ/LPS-treated macrophages, while SHP-2 co-precipitated with SIRPα under IL-4 stimulation (Fig. 2E).

### Src kinase-activated Btk phosphorylates SIRPα under IL-4/13 stimulation

Given that Lyn, a Src family tyrosine kinase, has been shown to phosphorylate ITIMs in different receptors including SIRPα (*23, 24*), we questioned the role of Lyn in IL-4/13-induced SIRPα phosphorylation. A Lyn-specific inhibitor Bafetinib (also named INNO-406) was tested; however, it showed only slight inhibition on IL-4/13-induced SIRPα phosphorylation (Fig. 3A). Instead, Bafetinib completely eliminated the low-level SIRPα phosphorylation induced by CD47 ligation in the absence of IL-4/13. These results suggest that Lyn was not responsible for the robust SIRPα phosphorylation under IL-4/13 stimulation. Instead, Lyn appeared constitutively active with a low phosphorylation capacity. To identify the tyrosine kinase(s) activated by IL-4/13, we tested pharmacological inhibitors targeting other kinases and found that the Src family kinase (SFK) inhibitor PP1, but not PP2, and the Btk inhibitor LFM-A13, but not its non-inhibitory analog LFM-A11 (Fig. 3D), strongly inhibited SIRPα phosphorylation and SIRPα association with SHP-2 by IL-4 (Fig. 3, B-C). To validate that Btk was the kinase phosphorylating SIRPα, another Btk-specific inhibitor, Ibrutinib, was used (*25*). As with LFM-A13, Ibrutinib dose-dependently inhibited SIRPα phosphorylation and association with SHP-2 in an IL-4/13-dependent manner (Fig. 3D). Additional experiments confirmed that IL-4 and IL-13 activated Btk, which had increased phosphorylation at Tyr^223^ (Fig. 3E), a result consistent with reports by others (*26, 27*). Both Btk inhibitors were found to deplete Btk phosphorylation (Fig. 3F). Furthermore, inhibition of SFK by PP1 prevented Btk phosphorylation (Fig. 3F), suggesting that certain PP1-sensitive SFK mediated Btk activation under IL-4/13 stimulation (depicted in Fig. 3G).

**Figure 3.**
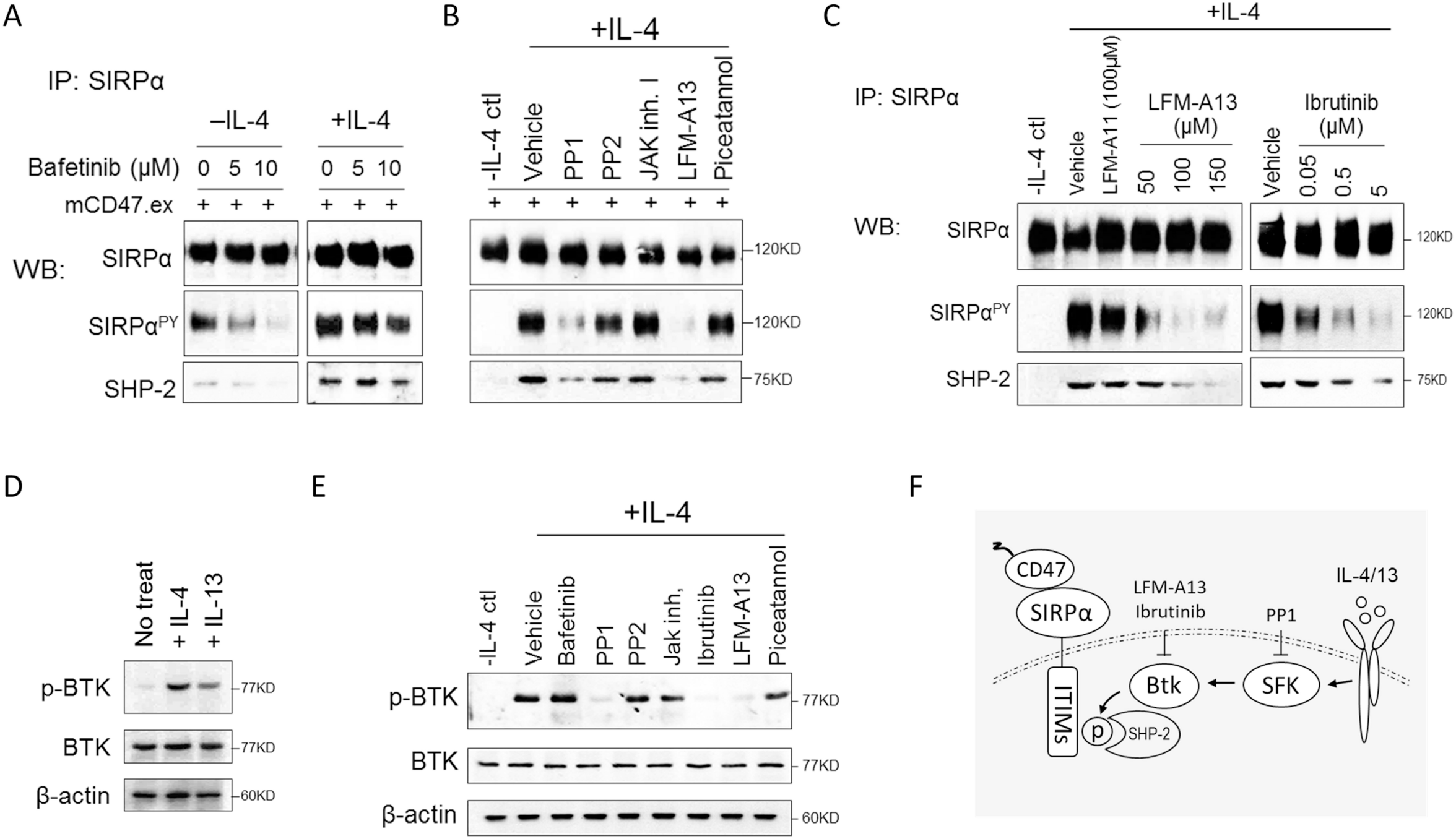
Src kinase-activated Btk phosphorylates SIRPα under IL-4/13 stimulation. Macrophages were activated by IL-4 (5ng/ml) in the presence of mCD47.ex for 20min to initiate downstream signaling of JAK, followed by inhibition of different tyrosine kinases using inhibitors. **(A)** Does-dependent effect of Bafetinib on SIRPα phosphorylation and its binding to SHP-2 in mCD47.ex and/or IL-4 (5ng/ml)-treated macrophages. **(B)** Effect of various tyrosine kinase inhibitors on SIRPα phosphorylation and SHP-2 association in IL-4 (5ng/ml)-activated macrophages. Data are presented as mean ± SEM of at least three independent experiments (n ≥ 3). Statistical significance was determined by Student’s t test where ***p < 0.001. **(C)** Dose-dependent effect of Btk inhibitor LFM-A13 or Ibrutinib on SIRPα phosphorylation and SHP-2 recruitment in IL-4-treated macrophages. Inactive LFM-A11 served as a negative control. **(D)** The phosphorylation of Btk after IL-4 (5ng/ml) or IL-13 (5ng/ml) treatment for 30 min. **(E)** Effect of different tyrosine kinase inhibitors on the phosphorylation of Btk in IL-4 activated macrophages. **(F)** Schematic depiction of the role of Btk in SIRPα phosphorylation and SHP-2 recruitment during IL-4-stimulated macrophage activation. All blots represent at least three independent experiments (n ≥ 3).

### Phosphorylated SIRPα sequesters SHP-2 to disinhibit IL-4/13 signaling

Given the finding that IL-4/13 stimulation induces SIRPα phosphorylation and recruitment of SHP-2, which coincidentally also binds to IL-4/13 receptors and inhibits IL-4/13 signaling, we developed a hypothesis postulating that the SIRPα-to-SHP-2 recruitment acts as a ‘disinhibition’ mechanism that, by SIRPα sequestering SHP-2, physically restricts SHP-2 from accessing the IL-4/13 receptors and thereby enhances cytokine-mediated signal transduction. This hypothesis agreed with the observation that SIRPα promoted macrophage alternative activation. To further test this hypothesis, we examined IL-4 and IL-13 receptors for phosphorylation and association with SHP-2 following cytokine stimulation in the presence or absence of SIRPα-mediated regulation (WT ± mCD47.ex or Sirpα^−/−^). As expected, IL-4 and IL-13 stimulation dose-dependently induced tyrosine phosphorylation of their heterodimeric receptor chains, IL-4Rα and γC or IL-13Rα1; however, only IL-4Rα exhibited varied phosphorylation correlated with SIRPα-mediated regulation. As shown (Fig. 4A), under IL-4/13 stimulation, WT macrophages ligated by CD47 (+ mCD47.ex) exhibited enhanced IL-4Rα phosphorylation compared to macrophages without CD47 ligation or depleted of SIRPα (Sirpα^−/−^), the latter indeed displaying diminished IL-4Rα phosphorylation even under high-dose cytokine stimulation. The level of IL-4Rα phosphorylation was also found to correlate with receptor-mediated downstream signaling, showing significantly increased or reduced activation of STAT6 and Akt1/2 and association of Gab2 with IL-4Rα. The co-receptor γC or IL-13Rα1, on the other hand, displayed minimal phosphorylation variations in the presence or absence of SIRPα-mediated regulation. These results suggest that the phosphorylation of IL-4Rα, but not its co-receptor γC or IL-13Rα1, is affected by SIRPα and impacts downstream signaling. This agrees with previous reports showing that IL-4Rα is the principal signaling chain whose phosphorylation provides docking of downstream molecules (*22*).

**Figure 4.**
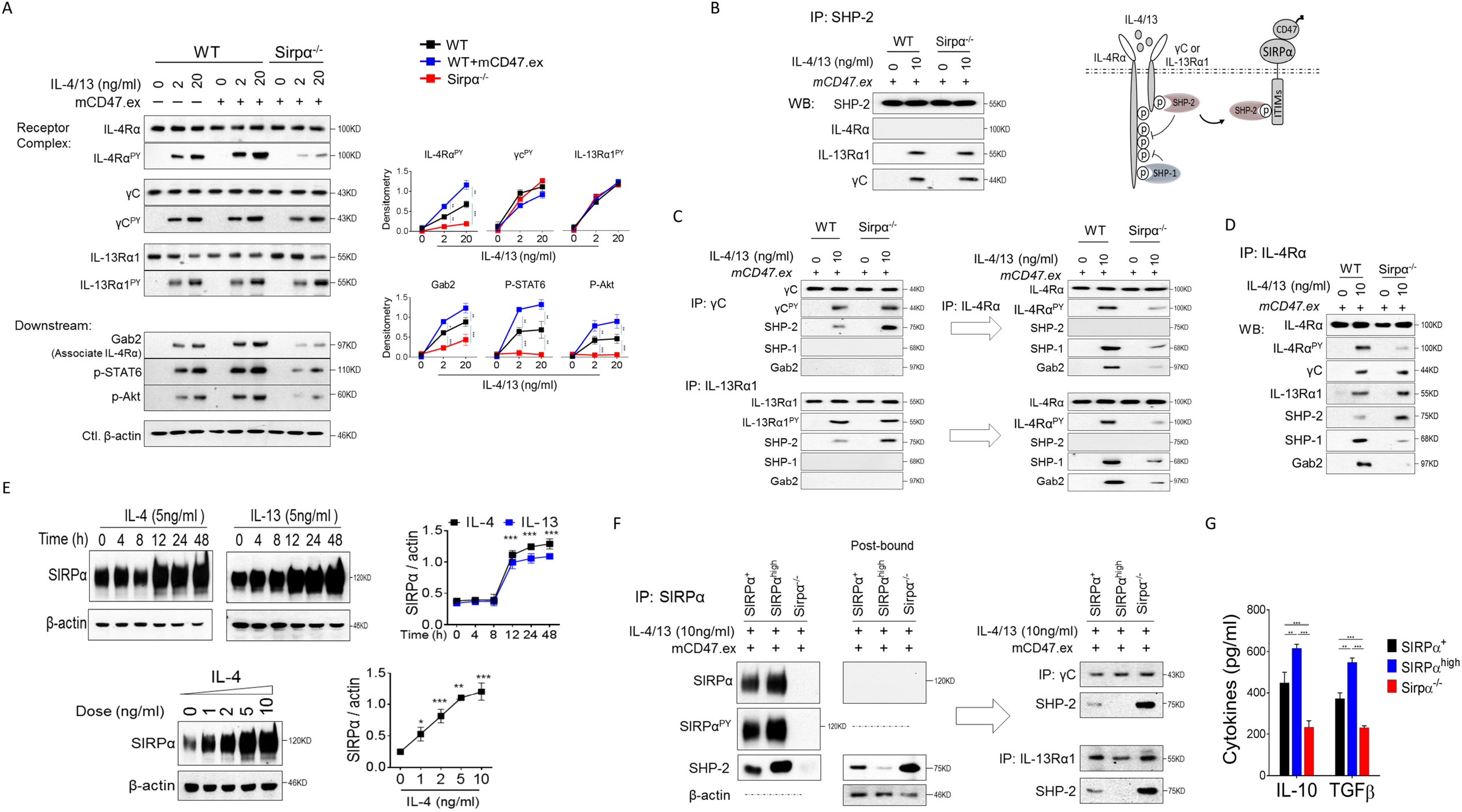
Phosphorylated SIRPα sequesters SHP-2 to disinhibit IL-4/13 signaling. **(A)** WT and Sirpα^−/−^ BMDM were treated with IL-4/13 (2 or 20ng/ml each) in the presence or absence of mCD47.ex for 20min. The phosphorylation of IL-4Rα, γ chain or IL-13Rα1 was detected by IP of IL-4Rα, γ chain or IL-13Rα1, respectively. The Gab2 associated with IL-4Rα and the phosphorylation of STAT6 and Akt in WT and Sirpα^−/−^ BMDM was also detected. **(B)** The binding of SHP-2 to IL-4Rα, γ chain or IL-13Rα1 in WT and Sirpα^−/−^ BMDM after IL-4/13 (10ng/ml each) treatment was detected by IP of SHP-2. **(C)** Sequential IP of γ chain, IL-13Rα1 and IL-4Rα from total cell lysate followed by WB of γ chain^PY^, IL-13Rα1^PY^, IL-4Rα^PY^, SHP-1, SHP-2 and Gab2. **(D)** The IL-4/13 receptor complexes were analyzed by IP of IL-4Rα. **(E)** Time course of SIRPα expression in WT BMDM after IL-4 (5ng/ml) or IL-13 (5ng/ml) and dose-dependent regulation of SIRPα by IL-4 after 24h. **(F)** WT, SIRPα^high^ or Sirpα^−/−^ BMDM were treated with IL-4/13 (10ng/ml each) and the association of SHP-2 with SIRPα, γ chain or IL-13Rα1 were assessed by IP of SIRPα, γ chain or IL-13Rα1, respectively. **(G)** The production of IL-10 and TGFβ from WT (SIRPα^+^), SIRPα^high^ and Sirpα^−/−^ macrophages in response to IL-4/13 (10ng/ml) after 24h. Data were presented as mean ± SEM of at least three independent experiments (n ≥ 3). Statistical significance was determined by Student’s t test (**A**, **E**, **G**) or where *p < 0.05, **p < 0.01 and ***p < 0.001. All blots are representative of at least three independent experiments.

IP of SHP-2 and detection of its co-associated cytokine receptors found that SHP-2 associated with γC and IL-13Rα1, but not IL-4Rα, in IL-4/13-stimulated macrophages (Fig.4B). Vice versa, IP of γC or IL-13Rα1 detected SHP-2 (Fig.4C). Given that the antibodies used to precipitate γC and IL-13Rα1 interrupt IL-4/13 receptor complexes and thus do not co-precipitate with IL-4Rα, the post-bound lysates depleted of γC or IL-13Rα1 were subsequently used for IP of IL-4Rα; however, these experiments failed to detect SHP-2 association. On the other hand, IP of IL-4Rα with an antibody that maintains IL-4Rα-γC/IL-13Rα1 receptor complexes co-precipitated IL-4Rα, γC and/or IL-13Rα1, and SHP-2 (Fig.4D). Comparing different macrophages (Fig.4 C-D) revealed that SHP-2 poorly bound to γC or IL-13Rα1 in WT macrophages with elevated SIRPα cytoplasmic phosphorylation (+ mCD47.ex) and that this reduction was associated with increased IL-4Rα phosphorylation. In stark contrast, SHP-2 highly associated with γC or IL-13Rα1 in Sirpα^−/−^ macrophages upon cytokine stimulation, and this increased γC/IL-13Rα1 association correlated with substantially reduced phosphorylation in IL-4Rα, leading to diminished downstream signaling. Thus, the extent to which SHP-2 bound to γC or IL-13Rα1 was inversely correlated with IL-4Rα phosphorylation and IL-4Rα-mediated downstream signal activation. Together, these results strongly supported our hypothesis that SIRPα sequestering SHP-2 serves to dis-inhibit IL-4/13 signaling, and also revealed that SHP-2 docks to γC or IL-13Rα1 while mediating dephosphorylation in IL-4Rα, which controls downstream signaling (depicted in Fig. 4B).

In tangential experiments, we observed that IL-4/13 stimulation significantly increased SIRPα expression in macrophages (Fig. 4 E). However, this response occurred slower (requiring > 8h) than IL-4/13-driven induction of immunosuppressive cytokines and other phenotypic markers (generally < 6h). Nevertheless, increased SIRPα expression (SIRPα^high^) enhanced SIRPα-to-SHP-2 sequestration to a greater extent, nearly depleting all SHP-2 available to inhibit the IL-4/13 signaling, thereby further sensitizing macrophages for alternative activation (Fig. 4F). Consequently, SIRPα^high^ macrophages, especially in the presence of CD47 ligation, produced much higher levels of IL-10 and TGFβ upon IL-4/13 stimulation (Fig. 4G). Thus, IL-4/13 signaling and SIRPα concertedly form a positive-feedback loop wherein: IL-4/13 increases SIRPα expression and thereby enhances SIRPα-mediated disinhibition through sequestration of SHP-2, a regulation that then bolsters IL-4/13 signaling to promote macrophage alternative activation.

### SIRPα directly regulates neither SHP-1 nor integrin under IL-4/13 signaling

The co-IP experiments found that SHP-1 also bound to IL-4/13 receptor complexes (Fig. 4C) and this finds was consistent with studies by the Chatila group (*7, 28*). Differing from SHP-2 that bound to γC or IL-13Rα1, SHP-1 bound to IL-4Rα and was in a manner that positively correlated with the level of IL-4Rα phosphorylation. Though SIRPα did not recruit SHP-1 under IL-4/13, it affected SHP-1-to-IL-4Rα binding through regulating SHP-2-mediated dephosphorylation of IL-4Rα. As shown (Fig. 4C), increased SIRPα sequestration of SHP-2 (WT +mCD47.ex) elevated not only IL-4Rα phosphorylation but also IL-4Rα-SHP-1 association, whereas SIRPα deficiency led to strong SHP-2-mediated dephosphorylation of IL-4Rα, diminishing SHP-1-to-IL-4Rα binding. While the manner of SHP-1-IL-4Rα association paralleled IL-4Rα phosphorylation and IL-4Rα-mediated downstream signaling activation, these results did not imply that SHP-1 positively regulates IL-4/13 signaling. Further studies employing phosphatase inhibitors confirmed that SHP-1, also SHP-2, inhibited IL-4/13 signaling and subsequent cytokine production (Fig. 5A). Inhibitors targeting either SHP-1 (TPI-1 and PTP-I) or SHP-2 (PHPS1 and SHP099) were tested, all of which dose-dependently increased IL-4/13-induced IL-10 and TGFβ production. As expected, SHP-2 inhibitors profoundly affected Sirpα^−/−^ macrophages but not SIRPα^high^ macrophages, as in the latter SHP-2 activity was largely negated by SIRPα. Meanwhile, inhibition of SHP-1 further increased the already high levels of IL-10 and TGFβ in SIRPα^high^ macrophages, but was less impactful toward Sirpα^−/−^ macrophages in which IL-4/13 signaling was dominantly controlled by SHP-2. Combining SHP-1 and SHP-2 inhibitors, however, drastically augmented cytokine production in all macrophages, surpassing that which was achieved by either inhibition.

**Figure 5.**
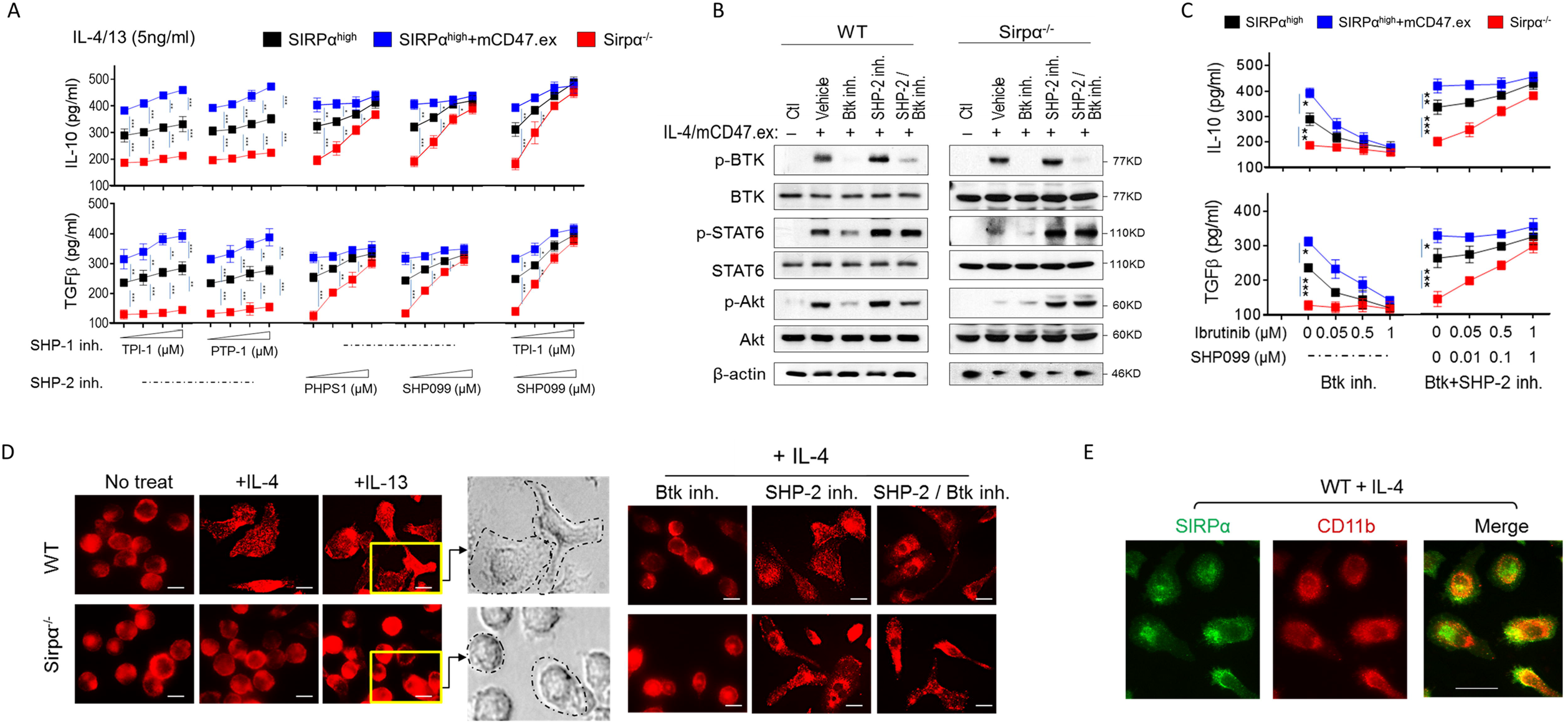
SHP-2 inhibition restores IL-4 signaling in SIRPα-deficient macrophages whereas Btk inhibition diminishes IL-4 signals in WT macrophages. **(A)** Effect of dose-dependent inhibition of SHP-1 (TPI-1 (0.005, 0.05 and 0.5μM) or PTP-I (1, 10 and 100 μM)), SHP-2 (PHS1 (0.5, 5 and 50 μM) or SHP099 (0.01, 0.1 and 1μM)) or both on the production of IL-10 (upper) and TGFβ (lower) in macrophages treated by IL-4/13 (5ng/ml, each) for 16h. **(B)** Effect of Btk inhibitor, SHP-2 inhibitor or the combination on IL-4-induced phosphorylation of STAT6 and Akt in WT and Sirpα^−/−^ macrophages. **(C)** Effect of dose-dependent inhibition of SHP-2 (SHP099) or Btk (Ibrutinib) on the production of IL-10 (upper) and TGFβ (lower) in macrophages treated by IL-4/13 (5ng/ml, each) for 16h. Data were presented as mean ± SEM of at least three independent experiments (n ≥ 3). Statistical significance was determined by Student’s t test (**A**, **C**) where *p < 0.05, **p < 0.01 and ***p < 0.001. **(D)** Immunostaining of CD11b on WT and Sirpα^−/−^ PEM in response to IL-4 (5ng/ml) or IL-13 (5ng/ml) in the presence of Btk inhibitor, SHP-2 inhibitor or both. Scale bar = 10 μm. **(E)** Immunostaining of CD11b and SIRPα on WT PEM. Scale bar = 10 μm. All blots are representative of at least three independent experiments.

Employing the SHP-2 inhibitor SHP099 and the Btk inhibitor Ibrutinib, we further validated the IL-4/13→Btk→SIRPα-SHP-2→IL-4/13 regulatory axis. Supporting our previous findings, Ibrutinib inhibited Btk activation (p-Btk), which is integral to SIRPα phosphorylation and SHP-2 sequestration, and dampened cytokine signaling in IL-4-stimulated macrophages (i.e., reduced p-STAT6 and p-Akt). While SHP099 did not affect Btk, its inhibition of SHP-2 resulted in increased phosphorylation of STAT6 and Akt in WT and Sirpα^−/−^ macrophages alike. The combination of Ibrutinib and SHP099 resulted in signal transduction similar to employing SHP099 alone, confirming that Btk is upstream of SIRPα-SHP-2 sequestration (Fig. 5B). Assaying cytokine production confirmed these results, showing Ibrutinib alone dose-dependently reduced IL-10 and TGFβ in SIRPα^+/high^ macrophages by IL-4, whereas the combination of Ibrutinib and SHP099 led to increased cytokine production as that which occurred under SHP-2 inhibition by SHP099 (Fig. 5C).

As SIRPα was reported to regulate leukocyte function through its cooperation with integrin-mediated cell adhesion (*29*), we investigated whether SIRPα deficiency alters cytokine-induced macrophage adhesion/spreading and if this occurs through the β2 integrin CD11b/CD18. Stimulation of WT and Sirpα^−/−^ macrophages induced their adhesion to the matrix, but only WT macrophages displayed an activated, broad spreading and/or elongated morphology (Fig. 5D). Comparably, IL-4-stimulated Sirpα^−/−^ macrophages exhibited a much less outstretched morphology suggestive of insufficient activation. Further studies confirmed that this difference in morphology was not due to the absence of SIRPα, but rather was from SHP-2 activity-reduced IL-4 signaling, as inhibition of SHP-2 by SHP099 enhanced IL-4 signaling and restored Sirpα^−/−^ macrophage spreading. In contrast, impairing SIRPα-to-SHP-2 sequestration in WT macrophages by inhibition of Btk (Ibrutinib) attenuated their response to IL-4 and spreading, which was restored by concomitant inhibition of SHP-2 (Ibrutinib + SHP099) (Fig. 5D). Staining of SIRPα and CD11b revealed punctate patterns on WT macrophage surface, though these two proteins did not co-localize (Fig. 5E). Together, these data suggest that SIRPα unlikely interacts with β2 integrin, but affects macrophage adhesion/spreading through regulating cytokine signaling.

### SIRPα deficiency impairs post-colitis wound healing by reducing IL-10^+^ macrophages

Alternatively activated macrophages orchestrate protective functions under inflammatory conditions and are critical for inflammation resolution (*30, 31*). Given our finding that SIRPα promotes alternative activation, we questioned whether SIRPα deficiency in vivo would impair immunosuppressive macrophage polarization and undermine disease resolution. Experimental colitis was induced in WT and Sirpα^−/−^ mice by feeding animals with 1-3% dextran sodium sulfate (DSS)-containing water for 6 days, and afterward mice were given DSS-free water to allow recovery. Consistent with previous studies (*14, 32*), Sirpα^−/−^ mice compared to WT mice developed more severe colitis under the same dose of DSS treatment (D1-D6), displaying faster body weight loss and higher fecal concentrations of lipocalin 2 (Lcn-2, a useful clinical marker of colitis) (Fig. 6A). Moreover, Sirpα^−/−^ mice also displayed difficulty to recover after DSS removal. As shown, compared to WT mice that passed the disease apex by D9 and restored their body weight to > 95% by D14, Sirpα^−/−^ mice exhibited protracted colitis and did not begin to recover until D15, followed by a slow recovery that would not reach 90% of their initial body weight by D20-30. Examination of cytokines found increases in circulating IL-6 but not IL-17A in Sirpα^−/−^ mice (Fig. 6C), despite that the latter has been suggested to drive colitis in other studies (*32, 33*). Inspection of resected colon tissues confirmed that Sirpα^−/−^ mice experienced worse colitis and slow wound healing, which was associated with a paucity of IL-10^+^ macrophages but an abundance of IL-6^+^ macrophages and neutrophils (Ly6G^+^) along the mucosal lining after DSS removal (D12) (Fig. 6 B-D). Speculating that Sirpα^−/−^ macrophage inability to undergo alternative activation underpinned the failure of inflammation resolution, we prepared IL-4-induced alternatively activated macrophages (WT/SIRPα^high^ BMDM, GFP^+^) ex vivo and intravenously transfused them into colitic Sirpα^−/−^ mice. Full body bioluminescent imaging revealed that IL-4-treated BMDM readily infiltrate the inflamed intestines in Sirpα^−/−^ mice (Fig. 6E). Following two rounds of transfusion (D6, D9), Sirpα^−/−^ mice markedly improved the rate at which they recovered from acute colitis, displaying both rapid body weight increase and clearance of fecal Lcn-2 (Fig. 6F).

**Figure 6.**
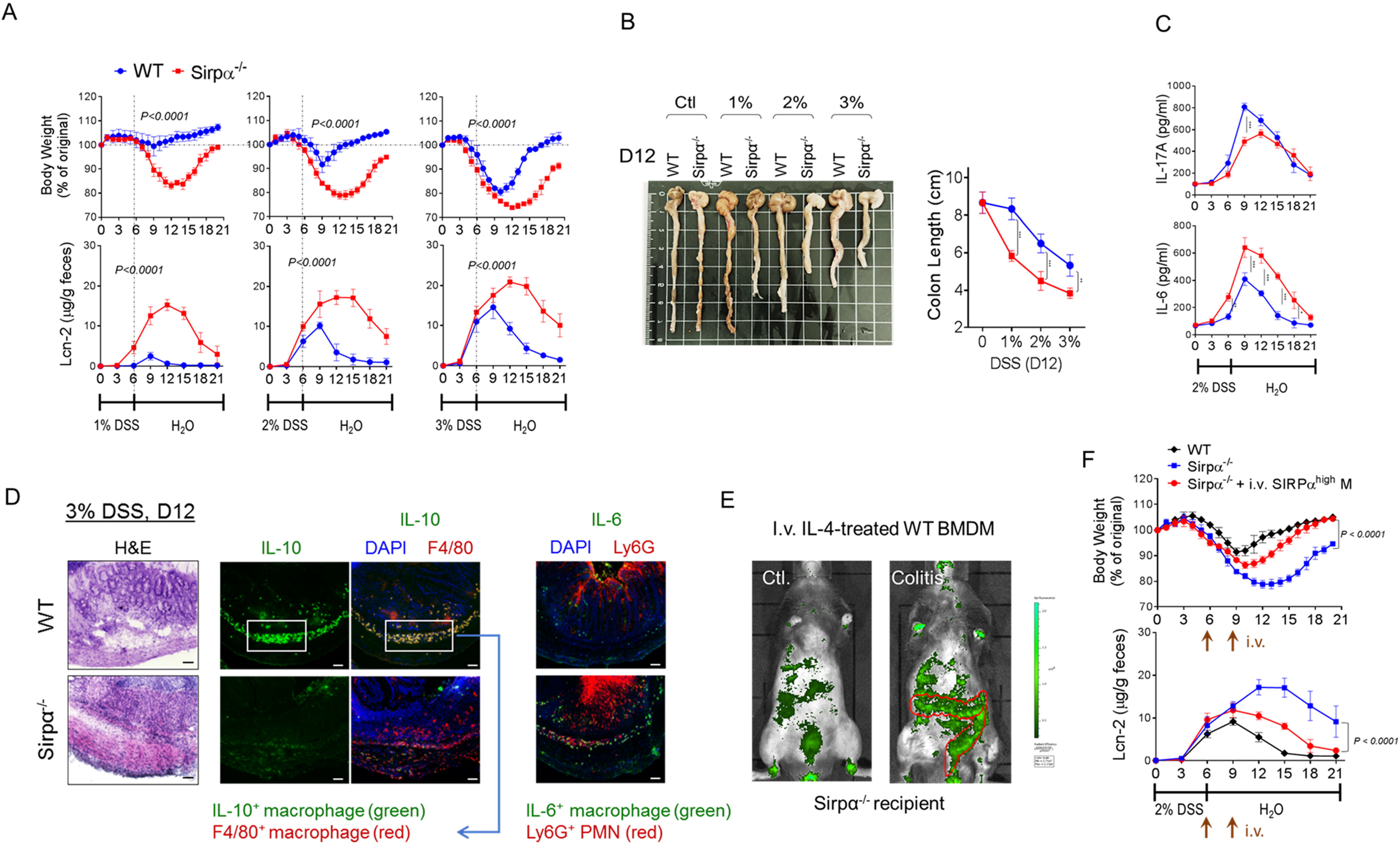
SIRPα deficiency impairs recovery from acute colitis in mice. **(A-D)** WT or Sirpα^−/−^ mice were treated with 1%, 2% or 3% DSS to induce acute colitis (D1-6), followed by allowing recovery (D6-21). Development of colitis was monitored by body weight loss and fecal Lcn-2 **(A)** and shortened intestines (D12, **B**). The levels of IL-17 and IL-6 in serum **(C)** and colon sections showing infiltration of IL-10- and IL-6-expressing macrophages (red and green, respectively) and neutrophil (Ly6G^+^, red) into intestines **(D)** were measured. Scale bar = 50μm. **(E-F)** 2% DSS-induced colitic Sirpα^−/−^ mice were intravenously administered with ex vivo IL-4/13-produced BMDM (SIRPα^high^ M, 1×107, GFP^+^) on D6 and D9 **(E)**; body weight change and fecal Lcn-2 were monitored **(F)**. Data in each panel represent at least three independent experiments with n ≥ 3 if applicable. Error bars are ± SEM. Statistical significance was determined by two-way ANOVA with Tukey’s multiple comparisons test (**A**, **F**) or Student’s t test (**B**, **C**), where *p < 0.05, **p < 0.01 and ***p < 0.001.

## Discussion

In addition to controlling macrophage phagocytosis of self-cells, SIRPα critically regulates macrophage responses to pro- and anti-inflammatory cues and phenotypic activation. Here, we show that macrophages, especially those with high SIRPα expression and extracellularly ligated by CD47, are significantly augmented for polarization by Th2 cytokines IL-4 and IL-13, exhibiting increased expressions of immunosuppressive cytokines IL-10 and TGFβ and alternative activation phenotypic markers Arg-1 and CD206. In contrast, macrophages deficient of SIRPα displayed an inability to undergo altenative activation due to much weakened responses to IL-4/13. Together, these studies ascertained that SIRPα promotes a macrophage anti-inflammatory phenotype that is conducive to tissue support and immunosuppression.

The mechanism by which SIRPα promotes IL-4/13-induced macrophage alternative activation revealed in this study was, however, unexpected. First, although SIRPα cytoplasmic ITIMs/ITSMs phosphorylation was expected to mediate the regulation, phosphorylation was triggered by cytokine stimulation but not CD47 extracellular binding. Indeed, IL-4/13 induces robust SIRPα phosphorylation through activating a SFK-sensitive tyrosine kinase Btk. The chain of these reactions, IL-4/13→SFK→Btk→SIRPα, occurs independently of CD47, albeit the latter ligating the SIRPα extracellular domain triggers SIRPα cytoplasmic structural change that facilitates the extent to which its tyrosine residues become phosphorylated. In addition to cytokine-activated Btk, Lyn also phosphorylates SIRPα cytoplasmic domain. However, our study revealed that Lyn plays a trivial role under cytokine stimulation, but its constitutive, low kinase activity is rather useful in maintaining macrophage homeostatic function once SIRPα is ligated by CD47.

Second, this mechanism of IL-4/13-induced SIRPα regulation functions like a loop of “self-authorization” that permits IL-4/13 receptors to proceed with signal transduction. Following IL-4/13 ligation, the authority for their receptors to propogate signaling is dependent on receptor tyrosine phosphorylation, which provide phospho-tyrosine (pY) docking sites for activation of downstream signal molecules. However, phosphorylated receptors recruit not only signaling molecules but also the pY-binding inhibitory molecules SHP-1 and SHP-2, which dephosphorylate IL-4/13 receptors and diminish signal transduction. Indeed, these phosphatase-mediated inhibitory activities are strong and largely control IL-4/13 signaling. As observed in Sirpα^−/−^ macrophages where SHP-2 is unleashed to dephosphorylate IL-4/13 receptors, > 80% of IL-4/13-mediated signaling was abated, resulting in ineffective macrophage alternative activation and IL-10 and TGFβ production. In contrast, restriction of SHP-2 by SIRPα enabled strong IL-4/13 signaling and high macrophage phenotypic expression of immuosuppressive cytokines. Thus, the SIRPα-mediated sequestration of SHP-2 endorses IL-4/13 signaling. As further revealed in this study, IL-4/13 signaling is in turn upregulating SIRPα expression (SIRPα^high^), thereby forming a self-reinforcing loop, IL-4/13→SIRPα→IL-4/13, that powerfully promotes macrophage polarization towards alternative activation.

Third, IL-4/13 induces phosphorylation of SIRPα ITIMs/ITSMs that controls only SHP-2, not SHP-1. Both corroborating and departing from previous work by Tachdjian et al. (*7*), we found that SHP-1 also inhibits IL-4/13 signaling. Different from SHP-2 that binds to the co-receptor γC or IL-13Rα1 and dephosphorylates the signaling chain IL-4Rα, SHP-1 directly docks to IL-4Rα (Y709). Though SIRPα does not physically recruit SHP-1, it impacts the degree of SHP-1 association to IL-4Rα by affecting SHP-2-mediated dephosphorylation of IL-4Rα. Presumably, when SHP-2 inhibition is strong, such as that in Sirpα^−/−^ macrophages, the role of SHP-1 is negligable, as the heavy dephosphorylation of IL-4Rα by SHP-2 depletes SHP-1 docking. Conversely, in SIRPα^high^ macrophages, where SHP-2 is restricted by SIRPα and, as such, IL-4Rα is less dephosphorylated, SHP-1 binding to IL-4Rα becomes prominent and its regulatory activity controls the IL-4/13 signaling strength.

The mechanism by which SIRPα recruits SHP-2 but not SHP-1 under IL-4/13 stimulation remains unclear. Two possibilities are conceivable: i) Btk phosphorylates SIRPα cytoplasmic tyrosines in a distinct pattern that confers binding affinity for only SHP-2 and not SHP-1. Though traditionally recognizing that the SIRPα cytoplasmic domain contains two ITIMs (IT*Y^433/436(human/murine)^*ADL and LT*Y^474/477^*ADL), it also has tyrosines residues forming two characteristic ITSMs (TE*Y^457/460^*ASI and SE*Y^500/501^*ASV). These ITIMs and ITSMs may be phosphorylated by different kinases under various activation conditions, forming phosphorylation patterns with differing affinities for either SHP-1 or SHP-2. Supporting this postulation, we observed that SIRPα mediates cytoplasmic recruitment of SHP-1 under proinflammatory stimulations (TLR ligands, IFNγ, IL-6, etc), while recruiting SHP-2 in response to IL-4, IL-13, IL-10 or TGFβ. Studies by others have also shown that ITIM and ITSM structures in different receptors are capable of binding to SHP-1 or/and SHP-2 (*34–37*). ii) Recruitment of SHP-1 or SHP-2 is dictated by alteration of the SH2 domain binding sites in these phosphatases following cytokine or other treatments. Despite their structural similarity, SHP-1 and SHP-2 are commonly found docking on different pY residues and perform distinct regulatory fiunctions. Between these two phosphatases, SHP-1 generally inhibits cellular functions after being recruited to pYs, like those in IL-4Rα, whereas the role of SHP-2 remains debated. Despite its capability of mediating inhibitory function – for example, when binding to γC or IL-13Rα1 and dephsohorylating IL-4Rα – SHP-2 ‘positively’ regulates cell signaling when recruited by SIRPα, as well as other molecules (*38, 39*). Perhaps, the differences between SHP-1 and SHP-2 are best examplified in animals deficient of these phosphatases (*40, 41*), although further investigation is clearly needed to completely understand these regulators.

Employing a murine colitis-recovery model, we here demonstrate that Sirpα^−/−^ mice not only develop more severe colitis due to enhanced macrophage and neutrophil functions (*42, 43*), but also exhibit greatly delayed resolution of inflammation and wound healing after DSS was removed, a condition associated with a lack of IL-10-expressing macrophages in the colon mucosa. Transfusing colitic Sirpα^−/−^ mice with SIRPα^high^ alternatively activated macrophages produced ex vivo redirected inflammatory condition towards resolution, leading to rapid wound healing and tissue recovery. After intravenous administration, these alternatively activated macrophages especially accumulated in the inflammatory intestines where they, instead of being reprogrammed by the ‘hot’ environment, directed an anti-inflammatory response that led to resolution of inflammation. Thus, depite that these experiments were designed to demonstrate the impact of SIRPα on macrophage alternative activation in vivo, they unexpectedly revealed a glimpse into the potential of using SIRPα^high^ macrophages as a cellular therapy for treating inflammatory diseases. Given that many of the current endeavors mitigate diseases, such as cytokine storm syndromes associated with COVID-19 and other infections, autoimmune inflammatory conditions and cancer, through immunotherapeutic modalities, leveraging the immune system by regulating SIRPα and directing macrophages to favor either pro-inflammatory or anti-inflammatory activation may provide a new avenue to rectify disease conditions and awaits further exploration.

## Materials and Methods

### Macrophages

Murine bone marrow-derived macrophages (BMDM) were produced by differentiation of bone marrow cells from WT and Sirpα^−/−^ mice (*14*) with macrophage colony stimulating factor (M-CSF)-conditioned RPMI-1640 medium with 10% FBS for 5 days (*14*). Murine peritoneal macrophages (PEM) were prepared by lavaging the peritoneal cavity with PBS followed by centrifugation and adherence to cell culture plates to remove non-adherent peritoneal cells (*14*). Human monocyte-derived macrophages (HMM) were produced by differentiating peripheral blood mononuclear cells (PBMC) in RPMI 1640 medium containing human M-CSF (10ng/mL, Biolegend) and 10% human platelets-free plasma for 5-7 days (*42, 44*). To prevent unwanted macrophage activation towards proinflammatory polarization, a caveat often caused by undetectable contamination, dust or unknown TLR agonists in the culture, low dosage IL-10 (2ng/ml) was sometimes added into the culture, which effectively maintained an inactive macrophage phenotype as indicated by assays detecting macrophage production of cytokines and macrophage metabolic states. Addition of IL-10 during HMM differentiation has also been described elsewhere (*45*). *Note:* Failure to control the macrophage phenotype during macrophage differentiation is the most common cause compromising studies of cell signaling and functionality induced by alternative activation stimuli. SIRPα^high^ macrophages were induced by IL-4 treatment (5 ng/ml, 16h) and were allowed to rest for >12h in non-IL-4-containing media prior to subsequent treatments.

### Soluble murine and human CD47 extracellular domains (mCD47.ex and hCD47.ex)

The AP-tag2 plasmid containing the extracellular domain of murine CD47 (mCD47.ex) and alkaline phosphatase (AP) was a generous gift of V. Narayanan (University of Pittsburgh School of Medicine) (*20*). This plasmid was then reconstructed by replacing mCD47.ex with the human CD47 extracellular domain (hCD47.ex). After transfecting COS cells, recombinant mCD47.ex and hCD47.ex fusion proteins were produced, affinity purified and stored in PBS as previously described (*11, 19–21*). Prior to use, the binding capacities of mCD47.ex or hCD47.ex with SIRPα were confirmed by their binding to murine or human SIRPα extracellular domain fusion protein, mSIRPα.ex-Fc or hSIRPα.ex-Fc, respectively, as well as adhesion to macrophages in a CD47-SIRPα-dependent manner (*21*).

### Macrophage alternative activation

IL-4 or IL-13 (Biolegend or PeproTech) was added into macrophage cultures at various concentrations to induce the M2/M2a phenotype in the presence or absence of a soluble CD47 extracellular domain (mCD47.ex or hCD47.ex). At different time points (0, 4, 10, 16 and 24h), cell-free supernatants were collected followed by ELISA to detect IL-10 and TGFβ using capture and detecting antibodies (Biolegend). Other M2 phenotype characteristics including the cell surface expressions of CD206, intracellular arginase (Arg-1) expression, and mRNA levels of Arg1, Ym1, Msr2 and Fizz1, were detected by a PE-conjugated anti-CD206 antibody (Biolegend) for flow cytometry, an anti-Arg-1 antibody (Santa Cruz Biotechnology) for western blot (WB) (*46, 47*), as well as oligonucleotide primers for real-time RT-PCR. For the last set of experiments, PCR primers were: Arg1, 5’-ctccaagccaaagtccttagag and 5’-aggagctgtcattagggacatc; Ym1, 5’-agaagggagtttcaaacctggt and 5’-gtcttgctcatgtgtgtaagtga; Msr2, 5’-aaagaaagcccgagtccc and 5’-tgcccaaggagatagcaaga; Fizz1, 5’-ggatgccaactttgaatagg and 5’-cttcgttacagtggagggat; GAPDH, 5’-tgaagcaggcatctgaggg and 5’-cgaaggtggaagagtgggag (*48*). In other experiments, macrophages were treated with IFNγ (20ng/ml), LPS (100ng/ml), IL-10 (20ng/ml) or TGFβ (20ng/ml) to induce different phenotypes in the presence or absence of mCD47.ex.

### Culture of bone marrow-derived dendritic cells (BMDCs)

Bone marrow-derived dendritic cells were induced by GM-CSF (20ng/ml) as previously described (*49*). On day 6, floating or lightly adherent cells were harvested by gentlely washing with PBS and then pooled for subsequent experiments. After 10 days induction, the cells were ready and treated with IL-4 for further analysis.

### Immunoprecipitation (IP) and WB

Murine macrophages were briefly treated with freshly prepared pervanadate (2mM, 5 min, 37°C) followed by lysis in an ice-cold buffer containing 25mM Tris-HCl, pH 7.4, 150mM NaCl, 1% Triton X-100, protease inhibitors (Protease Inhibitor Cocktail, Sigma-Aldrich), phosphatase inhibitors (Phosphatase Inhibitor Cocktail 1 and 2, Sigma), 3mM PMSF and 2mM pervanadate. After centrifugation (12,000rpm, 5 min), the clear lysates were collected. For IP, antibodies against SIRPα (clone P84, Biolegend), SHP-1, SHP-2, IL-4Rα, γ chain or IL-13Rα1 (all from Santa Cruz Biotechnology) were added at 5μg/ml followed by precipitation with protein A/G-Sepharose (4h, 4°C). After washing, the beads were heated at 95°C (5min) in SDS-PAGE sample buffer. After electrophoresis in acrylamide gels, proteins were transferred onto nitrocellulose, followed by blocking with 5% BSA and detection for SIRPα, SHP-1, SHP-2, IL-4Rα, γ chain or IL-13Rα1 using the same antibodies. The antibody for IP IL-4Rα could maintain the binding of IL-4Rα with γ chain or IL-13Rα1, whereas antibodies against γ chain or IL-13Rα1 caused the dissociation of IL-4Rα with γC or IL-13Rα1. Protein tyrosine phosphorylation was detected by PY20 (Biolegend) or 4G10 (Sigma), and phospho-SHP-2 was detected with an anti-phospho-SHP-2 (Tyr542) antibody (Cell Signaling Technology). Alternatively, macrophage lysates prior to IP or post-bound of IP were applied to SDS-PAGE and WB to detect the same proteins, as well as signal transduction molecules including Gab2, STAT6 and phospho-STAT6 (Tyr641), Akt1 and phospho-Akt1 (Ser473), Akt2 and phospho-Akt2 (Ser474), Akt and phospho-Akt (Ser473), Btk and phospho-Btk (Tyr223) (all antibody reagents from Cell Signaling Technology). The densitometry was done using the NIH software Image J.

### Inhibitor treatment

To inhibit SHP-2, specific inhibitors PHPS1 (Cayman Chemical, IC50 = 2.1μM) or SHP099 (Cayman Chemical, IC50 = 71nM) were used to treat macrophages prior to IL-4-induced alternative activation (*8*). To inhibit SHP-1, specific inhibitors TPI-1 (Cayman Chemical, IC50 = 40nM) or PTP Inhibitor I (PTP-I, Cayman Chemical, Ki = 43μM) were used to treat macrophages before IL-4 induction. To inhibit Lyn, a Lyn-specific inhibitor Bafetinib (5 or 10μM, Cayman Chemical, IC50 = 19nM)) was used to treat macrophages with or without ligation by mCD47.ex and/or treatment by IL-4 (*23*). To identify other tyrosine kinase(s) that phosphorylate SIRPα, the IL-4-stimulated macrophages were treated with the Src family kinase inhibitor PP1 (40μM, IC50 = 170nM) or PP2 (20μM, IC50 = 100nM), the pan-JAK inhibitor JAK inhibitor I (100nM, IC50 = 15nM), the Btk inhibitor Ibrutinib (0.05, 0.5 and 5μM, IC50 = 0.5nM) or LFM-A13 (50, 100 and 150μM, IC50 = 2.5μM), or Syk inhibitor piceatannol (40μM, IC50 = 10μM) (all from Cayman Chemical) for 20min, prior to treating with pervanadate and cell lysis (*50–52*). Subsequent IP and WB were performed to detect SIRPα phosphorylation and SHP-2 association. An LFM-A13 non-inhibitory analogue, LFM-A11 (150μM), was used as a control.

### DSS-induced colitis

To induce colitis, WT mice and Sirpα^−/−^ mice were treated with 1% DSS, 2% DSS or 3% DSS in drinking water for 6 days, followed by removal of DSS to allow recovery. To detect lipocalin 2 (Lcn-2) in feces, fecal samples were prepared in PBS with 0.1% Tween 20 (100 mg/ml) and were centrifuged to obtain clear supernatants, in which Lcn-2 levels were assayed using a murine Lcn-2 ELISA kit (R&D Systems). For macrophage adoptive transfer, GFP+ WT BMDM were treated with IL-4 (5ng/ml) for 16h. The Sirpα^−/−^ mice were intravenously transfused with IL-4 activated BMDM (1×107) 2 h after removal of DSS (D6) at three day intervals for a total of two treatments.

### Immunofluorescent tissue staining

DSS-treated mice were euthanized followed by harvesting of intestines. Intestines frozen in Tissue-Tek OCT were cryosectioned to 5-10μm slides, which were then fixed in ethanol and blocked with PBS containing 1% BSA (Sigma). H&E staining was performed to determine disease conditions. For immunofluorescence staining, slides were stained with rat anti-mouse IL-10, rat anti-mouse F4/80, rat anti-mouse IL-6 or rat anti-mouse Ly6G (all from Biolegend), followed by fluorescence-conjugated secondary antibodies. After washing, slides were mounted with DAPI (Invitrogen) and analyzed by fluorescent microscopy.

### Statistical analysis

All results represent at least three independent experiments. Quantitative data were analyzed in GraphPad Prism and are presented as means ± SEM, and differences were considered to be statistically significant when p < 0.05 by the Student’s t tests for paired samples. For multiple pairwise comparisons between samples and a shared control, analysis of two-way ANOVA with Tukey’s post hoc test was performed.

## Acknowledgments

This work was supported, in part, by the NIH grants AI106839 and CA241271, the Biolocity Innovation program and a venture development grant from the Georgia Research Alliance (GRA).

## Author contributions

Y.L. initiated the study and oversaw all data acquisition and analysis. L.S. performed all the key experiments with critical help from Z.B., K.K., S.C. and C.O.. L.S., K.K. and Y.L. wrote the manuscript.

## References

1. A. Shapouri-Moghaddam, S. Mohammadian, H. Vazini, M. Taghadosi, S. A. Esmaeili, F. Mardani, B. Seifi, A. Mohammadi, J. T. Afshari, A. Sahebkar, Macrophage plasticity, polarization, and function in health and disease. J. Cell. Physiol. 233, 6425–6440 (2018).

2. P. J. Murray, J. E. Allen, S. K. Biswas, E. A. Fisher, D. W. Gilroy, S. Goerdt, S. Gordon, J. A. Hamilton, L. B. Ivashkiv, T. Lawrence, M. Locati, A. Mantovani, F. O. Martinez, J. L. Mege, D. M. Mosser, G. Natoli, J. P. Saeij, J. L. Schultze, K. A. Shirey, A. Sica, J. Suttles, I. Udalova, J. A. van Ginderachter, S. N. Vogel, T. A. Wynn, Macrophage activation and polarization: nomenclature and experimental guidelines. Immunity 41, 14–20 (2014).

3. S. J. Van Dyken, R. M. Locksley, Interleukin-4- and interleukin-13-mediated alternatively activated macrophages: roles in homeostasis and disease. Annu. Rev. Immunol. 31, 317–343 (2013).

4. Z. Ul-Haq, S. Naz, M. A. Mesaik, Interleukin-4 receptor signaling and its binding mechanism: A therapeutic insight from inhibitors tool box. Cytokine Growth Factor Rev. 32, 3–15 (2016).

5. X. Guo, T. Li, Y. Xu, X. Xu, Z. Zhu, Y. Zhang, J. Xu, K. Xu, H. Cheng, X. Zhang, Y. Ke, Increased levels of Gab1 and Gab2 adaptor proteins skew interleukin-4 (IL-4) signaling toward M2 macrophage-driven pulmonary fibrosis in mice. J. Biol. Chem. 292, 14003–14015 (2017).

6. E. Vergadi, E. Ieronymaki, K. Lyroni, K. Vaporidi, C. Tsatsanis, Akt Signaling Pathway in Macrophage Activation and M1/M2 Polarization. J. Immunol. 198, 1006–1014 (2017).

7. R. Tachdjian, S. Al Khatib, A. Schwinglshackl, H. S. Kim, A. Chen, J. Blasioli, C. Mathias, H. Y. Kim, D. T. Umetsu, H. C. Oettgen, T. A. Chatila, In vivo regulation of the allergic response by the IL-4 receptor alpha chain immunoreceptor tyrosine-based inhibitory motif. J. Allergy Clin. Immunol. 125, 1128–1136 (2010).

8. B. Tao, W. Jin, J. Xu, Z. Liang, J. Yao, Y. Zhang, K. Wang, H. Cheng, X. Zhang, Y. Ke, Myeloid-specific disruption of tyrosine phosphatase Shp2 promotes alternative activation of macrophages and predisposes mice to pulmonary fibrosis. J. Immunol. 193, 2801–2811 (2014).

9. L. Zhao, J. Xia, T. Li, H. Zhou, W. Ouyang, Z. Hong, Y. Ke, J. Qian, F. Xu, Shp2 Deficiency Impairs the Inflammatory Response Against Haemophilus influenzae by Regulating Macrophage Polarization. The Journal of Infectious Diseases 214, 625–633 (2016).

10. S. S. Wang, Y. Y. Yao, H. X. Li, G. Zheng, S. Lu, W. B. Chen, Tumor-associated macrophages (TAMs) depend on Shp2 for their anti-tumor roles in colorectal cancer. Am. J. Cancer Res. 9, 1957-+ (2019).

11. Y. Liu, H. J. Buhring, K. Zen, S. L. Burst, F. J. Schnell, I. R. Williams, C. A. Parkos, Signal regulatory protein (SIRPalpha), a cellular ligand for CD47, regulates neutrophil transmigration. J. Biol. Chem. 277, 10028–10036 (2002).

12. Y. Liu, M. B. O’Connor, K. J. Mandell, K. Zen, A. Ullrich, H. J. Buhring, C. A. Parkos, Peptide-mediated inhibition of neutrophil transmigration by blocking CD47 interactions with signal regulatory protein alpha. J. Immunol. 172, 2578–2585 (2004).

13. K. Zen, Y. Guo, Z. Bian, Z. Lv, D. Zhu, H. Ohnishi, T. Matozaki, Y. Liu, Inflammation-induced proteolytic processing of the SIRPalpha cytoplasmic ITIM in neutrophils propagates a proinflammatory state. Nat Commun 4, 2436 (2013).

14. Z. Bian, L. Shi, Y. L. Guo, Z. Lv, C. Tang, S. Niu, A. Tremblay, M. Venkataramani, C. Culpepper, L. Li, Z. Zhou, A. Mansour, Y. Zhang, A. Gewirtz, K. Kidder, K. Zen, Y. Liu, Cd47-Sirpalpha interaction and IL-10 constrain inflammation-induced macrophage phagocytosis of healthy self-cells. Proc. Natl. Acad. Sci. U. S. A. 113, E5434–5443 (2016).

15. E. M. van Beek, J. A. Zarate, R. van Bruggen, K. Schornagel, A. T. Tool, T. Matozaki, G. Kraal, D. Roos, T. K. van den Berg, SIRPalpha controls the activity of the phagocyte NADPH oxidase by restricting the expression of gp91(phox). Cell Rep. 2, 748–755 (2012).

16. L. Shi, T. Bian, Y. Liu, Dual role of SIRP alpha in macrophage activation: inhibiting M1 while promoting M2 polarization via selectively activating SHP-1 and SHP-2 signal. J. Immunol. 198, (2017).

17. Y. Lin, J. L. Zhao, Q. J. Zheng, X. Jiang, J. Tian, S. Q. Liang, H. W. Guo, H. Y. Qin, Y. M. Liang, H. Han, Notch Signaling Modulates Macrophage Polarization and Phagocytosis Through Direct Suppression of Signal Regulatory Protein a Expression. Front. Immunol. 9, (2018).

18. T. Matozaki, Y. Murata, H. Okazawa, H. Ohnish, Functions and molecular mechanisms of the CD47-SIRP alpha signalling pathway. Trends Cell Biol. 19, 72–80 (2009).

19. Z. Lv, Z. Bian, L. Shi, S. Niu, B. Ha, A. Tremblay, L. Li, X. Zhang, J. Paluszynski, M. Liu, K. Zen, Y. Liu, Loss of Cell Surface CD47 Clustering Formation and Binding Avidity to SIRPalpha Facilitate Apoptotic Cell Clearance by Macrophages. J. Immunol. 195, 661–671 (2015).

20. P. H. Jiang, C. F. Lagenaur, V. Narayanan, Integrin-associated protein is a ligand for the P84 neural adhesion molecule. J. Biol. Chem. 274, 559–562 (1999).

21. Y. Liu, Q. Tong, Y. Zhou, H. W. Lee, J. J. Yang, H. J. Buhring, Y. T. Chen, B. Ha, C. X. Chen, Y. Yang, K. Zen, Functional elements on SIRPalpha IgV domain mediate cell surface binding to CD47. J. Mol. Biol. 365, 680–693 (2007).

22. S. M. McCormick, N. M. Heller, Commentary: IL-4 and IL-13 receptors and signaling. Cytokine 75, 38–50 (2015).

23. R. J. Cornall, J. G. Cyster, M. L. Hibbs, A. R. Dunn, K. L. Otipoby, E. A. Clark, C. C. Goodnow, Polygenic autoimmune traits: Lyn, CD22, and SHP-1 are limiting elements of a biochemical pathway regulating BCR signaling and selection. Immunity 8, 497–508 (1998).

24. H. Nishizumi, K. Horikawa, I. Mlinaric-Rascan, T. Yamamoto, A double-edged kinase Lyn: a positive and negative regulator for antigen receptor-mediated signals. J. Exp. Med. 187, 1343–1348 (1998).

25. T. Munir, J. R. Brown, S. O’Brien, J. C. Barrientos, P. M. Barr, N. M. Reddy, S. Coutre, C. S. Tam, S. P. Mulligan, U. Jaeger, T. J. Kipps, C. Moreno, M. Montillo, J. A. Burger, J. C. Byrd, P. Hillmen, S. Dai, A. Szoke, J. P. Dean, J. A. Woyach, Final analysis from RESONATE: Up to six years of follow-up on ibrutinib in patients with previously treated chronic lymphocytic leukemia or small lymphocytic lymphoma. Am. J. Hematol. 94, 1353–1363 (2019).

26. S. E. M. Herman, A. L. Gordon, E. Hertlein, A. Ramanunni, X. L. Zhang, S. Jaglowski, J. Flynn, J. Jones, K. A. Blum, J. J. Buggy, A. Hamdy, A. J. Johnson, J. C. Byrd, Bruton tyrosine kinase represents a promising therapeutic target for treatment of chronic lymphocytic leukemia and is effectively targeted by PCI-32765. Blood 117, 6287–6296 (2011).

27. A. J. Gunderson, M. M. Kaneda, T. Tsujikawa, A. V. Nguyen, N. I. Affara, B. Ruffell, S. Gorjestani, S. M. Liudahl, M. Truitt, P. Olson, G. Kim, D. Hanahan, M. A. Tempero, B. Sheppard, B. Irving, B. Y. Chang, J. A. Varner, L. M. Coussens, Bruton Tyrosine Kinase-Dependent Immune Cell Cross-talk Drives Pancreas Cancer. Cancer Discov. 6, 270–285 (2016).

28. G. K. K. Hershey, M. F. Friedrich, L. A. Esswein, M. L. Thomas, T. A. Chatila, The association of atopy with a gain-of-function mutation in the alpha subunit of the interleukin-4 receptor. N. Engl. J. Med. 337, 1720–1725 (1997).

29. F. J. Alenghat, Q. J. Baca, N. T. Rubin, L. I. Pao, T. Matozaki, C. A. Lowell, D. E. Golan, B. G. Neel, K. D. Swanson, Macrophages require Skap2 and Sirpalpha for integrin-stimulated cytoskeletal rearrangement. J. Cell Sci. 125, 5535–5545 (2012).

30. Y. Lin, X. Yang, W. Yue, X. Xu, B. Li, L. Zou, R. He, Chemerin aggravates DSS-induced colitis by suppressing M2 macrophage polarization. Cell. Mol. Immunol. 11, 355–366 (2014).

31. M. M. Hunter, A. Wang, K. S. Parhar, M. J. Johnston, N. Van Rooijen, P. L. Beck, D. M. McKay, In vitro-derived alternatively activated macrophages reduce colonic inflammation in mice. Gastroenterology 138, 1395–1405 (2010).

32. W. O’Connor, L. A. Zenewicz, R. A. Flavell, The dual nature of T(H)17 cells: shifting the focus to function. Nat. Immunol. 11, 471–476 (2010).

33. Z. Bian, Y. Guo, B. Ha, K. Zen, Y. Liu, Regulation of the inflammatory response: enhancing neutrophil infiltration under chronic inflammatory conditions. J. Immunol. 188, 844–853 (2012).

34. P. Scapini, S. Pereira, H. Zhang, C. A. Lowell, Multiple roles of Lyn kinase in myeloid cell signaling and function. Immunol. Rev. 228, 23–40 (2009).

35. B. E. Tourdot, M. K. Brenner, K. C. Keough, T. Holyst, P. J. Newman, D. K. Newman, Immunoreceptor Tyrosine-Based Inhibitory Motif (ITIM)-Mediated Inhibitory Signaling Is Regulated by Sequential Phosphorylation Mediated by Distinct Nonreceptor Tyrosine Kinases: A Case Study Involving PECAM-1. Biochemistry 52, 2597–2608 (2013).

36. Q. Zhang, W. B. Lee, J. S. Kang, L. K. Kim, Y. J. Kim, Integrin CD11b negatively regulates Mincle-induced signaling via the Lyn-SIRPalpha-SHP1 complex. Exp. Mol. Med. 50, e439 (2018).

37. J. M. Chemnitz, R. V. Parry, K. E. Nichols, C. H. June, J. L. Riley, SHP-1 and SHP-2 associate with immunoreceptor tyrosine-based switch motif of programmed death 1 upon primary human T cell stimulation, but only receptor ligation prevents T cell activation. J. Immunol. 173, 945–954 (2004).

38. X. J. Li, C. B. Goodwin, S. C. Nabinger, B. M. Richine, Z. Yang, H. Hanenberg, H. Ohnishi, T. Matozaki, G. S. Feng, R. J. Chan, Protein-tyrosine phosphatase Shp2 positively regulates macrophage oxidative burst. J. Biol. Chem. 290, 3894–3909 (2015).

39. C. Niogret, W. Birchmeier, G. Guarda, SHP-2 in Lymphocytes’ Cytokine and Inhibitory Receptor Signaling. Front. Immunol. 10, 2468 (2019).

40. Z. Z. Chong, K. Maiese, The Src homology 2 domain tyrosine phosphatases SHP-1 and SHP-2: diversified control of cell growth, inflammation, and injury. Histol. Histopathol. 22, 1251–1267 (2007).

41. U. Lorenz, SHP-1 and SHP-2 in T cells: two phosphatases functioning at many levels. Immunol. Rev. 228, 342–359 (2009).

42. K. Zen, Y. Guo, Z. Bian, Z. Lv, D. Zhu, H. Ohnishi, T. Matozaki, Y. Liu, Inflammation-induced proteolytic processing of the SIRPalpha cytoplasmic ITIM in neutrophils propagates a proinflammatory state. Nature Communications 4, 2436 (2013).

43. K. Kidder, Z. Bian, L. Shi, Y. Liu, Inflammation Unrestrained by SIRPalpha Induces Secondary Hemophagocytic Lymphohistiocytosis Independent of IFN-gamma. J. Immunol. 205, 2821–2833 (2020).

44. D. H. Zhu, C. Y. Pan, L. M. Li, Z. Bian, Z. Y. Lv, L. Shi, J. Zhang, D. H. Li, H. W. Gu, C. Y. Zhang, Y. Liu, K. Zen, MicroRNA-17/20a/106a modulate macrophage inflammatory responses through targeting signal-regulatory protein alpha. J. Allergy Clin. Immunol. 132, 426–+ (2013).

45. X. Jin, H. S. Kruth, Culture of Macrophage Colony-stimulating Factor Differentiated Human Monocyte-derived Macrophages. Journal of Visualized Experiments, (2016).

46. Z. Bian, A. M. Abdelaal, L. Shi, H. W. Liang, L. Q. Xiong, K. Kidder, M. Venkataramani, C. Culpepper, K. Zen, Y. Liu, Arginase-1 is neither constitutively expressed in nor required for myeloid-derived suppressor cell-mediated inhibition of T-cell proliferation. Eur. J. Immunol. 48, 1046–1058 (2018).

47. Z. Bian, L. Shi, M. Venkataramani, A. M. Abdelaal, C. Culpepper, K. Kidder, H. Liang, K. Zen, Y. Liu, Tumor conditions induce bone marrow expansion of granulocytic, but not monocytic, immunosuppressive leukocytes with increased CXCR2 expression in mice. Eur. J. Immunol. 48, 532–542 (2018).

48. X. Cai, Y. Yin, N. Li, D. Zhu, J. Zhang, C. Y. Zhang, K. Zen, Re-polarization of tumor-associated macrophages to pro-inflammatory M1 macrophages by microRNA-155. J. Mol. Cell. Biol. 4, 341–343 (2012).

49. A. Maroof, M. Penny, R. Kingston, C. Murray, S. Islam, P. A. Bedford, S. C. Knight, Interleukin-4 can induce interleukin-4 production in dendritic cells. Immunology 117, 271–279 (2006).

50. K. Zen, Y. Liu, Role of different protein tyrosine kinases in fMLP-induced neutrophil transmigration. Immunobiology 213, 13–23 (2008).

51. B. Lu, T. Nakamura, K. Inouye, J. Li, Y. Tang, P. Lundback, S. I. Valdes-Ferrer, P. S. Olofsson, T. Kalb, J. Roth, Y. Zou, H. Erlandsson-Harris, H. Yang, J. P. Ting, H. Wang, U. Andersson, D. J. Antoine, S. S. Chavan, G. S. Hotamisligil, K. J. Tracey, Novel role of PKR in inflammasome activation and HMGB1 release. Nature 488, 670–674 (2012).

52. A. Colado, M. Genoula, C. Cougoule, J. L. Marin Franco, M. B. Almejun, D. Risnik, D. Kviatcovsky, E. Podaza, E. E. Elias, F. Fuentes, I. Maridonneau-Parini, F. R. Bezares, H. Fernandez Grecco, M. Cabrejo, C. Jancic, M. D. C. Sasiain, M. Giordano, R. Gamberale, L. Balboa, M. Borge, Effect of the BTK inhibitor ibrutinib on macrophage- and gammadelta T cell-mediated response against Mycobacterium tuberculosis. Blood Cancer J. 8, 100 (2018).

53. P. C. Cook, L. H. Jones, S. J. Jenkins, T. A. Wynn, J. E. Allen, A. S. MacDonald, Alternatively activated dendritic cells regulate CD4(+) T-cell polarization in vitro and in vivo. Proc. Natl. Acad. Sci. U. S. A. 109, 9977–9982 (2012).

